# An Investigation into Patient-Specific 3D Printed Titanium Stents and the use of Etching to Overcome Selective Laser Melting Design Constraints

**DOI:** 10.1101/2021.03.07.434132

**Authors:** Orla M. McGee, Sam Geraghty, Celia Hughes, Parastoo Jamshidi, Damien P. Kenny, Moataz M. Attallah, Caitríona Lally

## Abstract

There is currently a clear clinical need in the area of stenting for paediatric patients, whereby currently commercially available adult stents are often required to be used off-label for paediatric patients resulting in less than optimal outcomes. The increasingly widespread use of CT and/or MR imaging for pre-surgical assessment, and the emergence of additive manufacturing processes such as 3D printing, could enable bespoke devices to be produced efficiently and cost-effectively. However, 3D printed metallic stents need to be self-supporting leading to limitations in the design of stents available through additive manufacturing. In this study we investigate the use of etching to overcome these design constraints and improve stent surface finish. Furthermore, using a combination of experimental bench testing and finite element methods we investigate how etching influences stent performance and using an inverse finite element approach the material properties of the printed and etched stents were calibrated and compared. Finally, using patient-specific finite element models the stent performance was tested to assess patient outcomes. Within this study, etching is confirmed as a means to create open-cell stent designs whilst conforming to additive manufacturing ‘rules’ and concomitantly improving stent surface finish. Additionally, the feasibility of using an in-vivo imaging to product development pipeline is demonstrated that enables patient-specific stents to be produced for varying anatomies to achieve optimum device performance.

**Figure 1.**
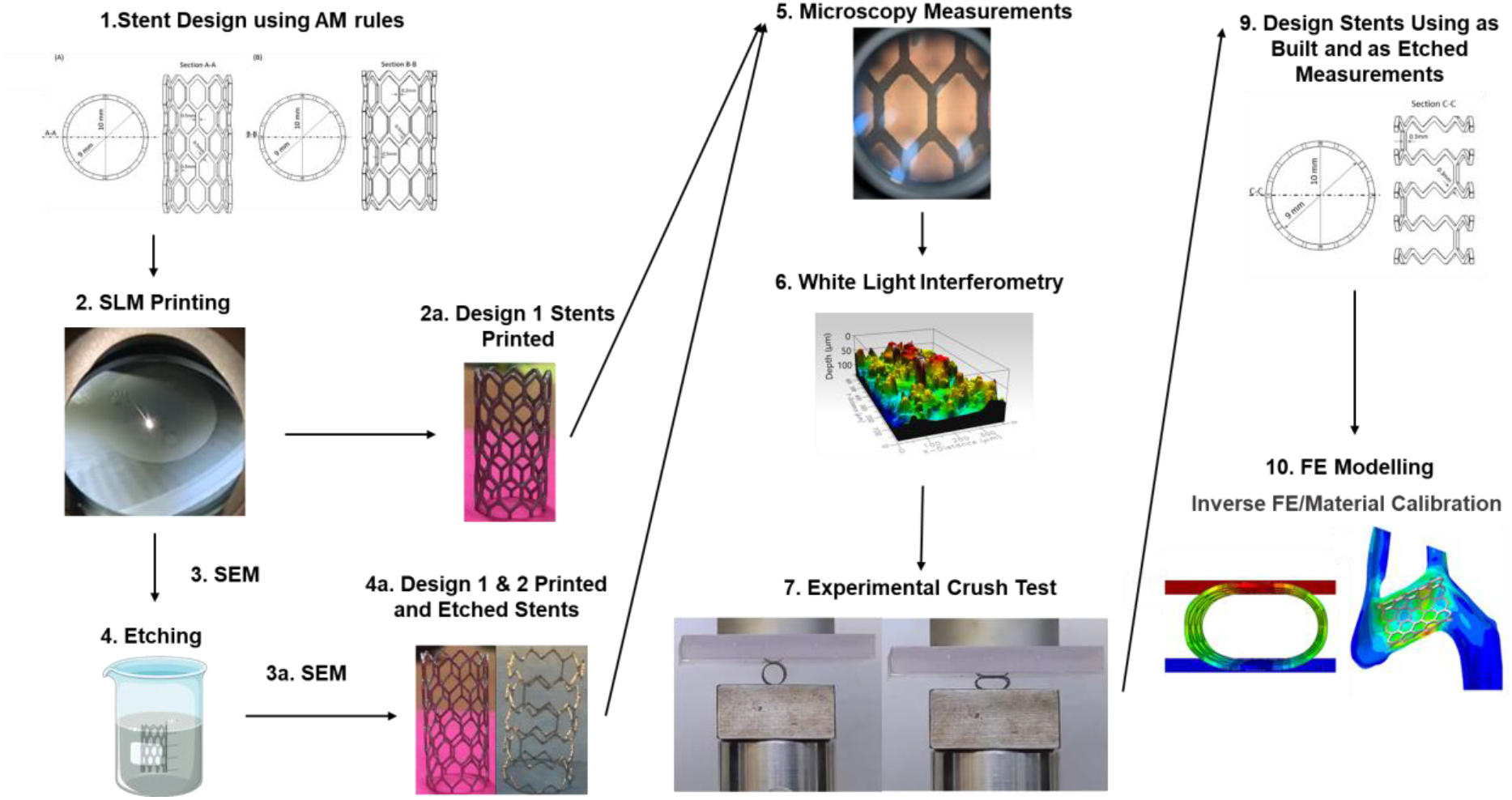
Graphical abstract.

## 1. Introduction

In the past 30 years, percutaneous therapies for the treatment of congenital heart disease have evolved immensely, however, with the limited investment from industry and support from regulatory bodies there is a lack of devices specifically designed for the treatment of congenital heart disease in paediatric patients [1]. This is further evidenced by the common off-label use of commercially available adult stents in the treatment of paediatric patients, often leading to less than optimal clinical outcomes [1,2]. A clear clinical need exists in this area of paediatrics and with the increasingly widespread use of Computed Tomography (CT) and/or Magnetic Resonance Imaging (MRI) for pre-surgical assessment and the emergence of Additive Manufacturing (AM) processes such as 3D printing, bespoke devices could be produced efficiently and cost-effectively to meet this need. This is particularly important in cases of congenital heart disease as children born with cardiovascular defects present a wide range of different anatomies and conditions that are often unique and complex [3,4].

AM capabilities have allowed for the design of components with reduced weight, improved mechanical performance and reduction in the cost of production [5]. AM techniques for medical applications, including biomedical implants, have been expanding rapidly and 3D printing of customised patient-specific device designs are expected to revolutionise future healthcare [6].

In recent years, research has been carried out into the development of 3D polymer stents [7–10]. Misra et al. developed a patient-specific polymer stent whereby 3D segmentation of medical images and in silico modelling were used to optimize the stent design [8]. However, with 99% of stents used in the treatment of coronary artery disease consisting of metal balloon-expandable stents [11] and the most widely deployed cardiovascular stents made of stainless steel, titanium and cobalt-chromium [5,12], there is a need for further research into the potential for AM of patient-specific metallic stents. Recently, 3D printed metallic stents have been developed in an attempt to eliminate the drawbacks of conventional stent manufacturing techniques where stents are laser cut from a tube [5,13]. These drawbacks include stent dimensions being dependent on available tubes with high production costs. Furthermore, using standard tube diameters can lead to the use of oversized stents inducing an inflammatory response when implanted [13–15]. Reduction in this inflammatory response reduces the likelihood of in-stent restenosis, the most significant long-term limitation of stenting [16]. Metallic stents produced via AM could overcome these challenges in both production and treatment leading the way to 3D printing bespoke and unique stent designs. This would be particularly useful for paediatric patients who require unique device designs. This paired with a relatively small patient cohort and challenges related to medical device approval results in little financial incentive in the development of devices for these patients [17].

Selective Laser Melting (SLM) is one AM method which has already been used to 3D print metallic stents [5,13,18]. SLM is a technique whereby successive layers of powder are selectively melted using a laser beam in order to generate a specified geometry [13]. A study by Demir & Previtali [5] investigated SLM of cobalt-chromium stents for cardiovascular stenting. That study outlined rules on creating a geometry that is self-supporting and negates the need for supports in the printing of the stent. This ensures there is no unnecessary damage to the stent in removing these supports post-printing. These rules state that struts at angles greater than 45° to the horizontal, overhangs less than 1mm, and gaps greater than 0.3mm should be incorporated into the print designs. However, these design rules lead to restrictions in the design of stents available through AM. Furthermore, the printing of SLM stents requires further investigation as limitations to printing on the micro-scale such as resolution, surface finish and layer bonding can impact the stent performance [19].

Finite Element (FE) modelling has been used over the past two decades to investigate cardiac mechanics, hemodynamics and device design [20]. In particular, patient-specific FE models have been used to investigate cardiovascular device performance, optimal sizing, placement and optimization of device design [3,4,21–23]. Li et al. previously used FE methods to optimize the design of coronary stents to reduce the stent dog-boning effect [23]. Rocatello et al. recently demonstrated the feasibility of using a patient-specific FE model to optimize the device design of a heart valve stent frame suggesting that such methods could be used prior to the first-in-human trials to develop improved devices [22].

The objective of this study was to determine the feasibility of generating patient-specific SLM stents and the use of etching to overcome design restrictions of the SLM technique. In this study, we used CT images of a 4-year old patient with a congenital heart disease known as aortic coarctation, requiring treatment with a stent. The patient’s geometry was reconstructed and the design of a patient-specific stent using SLM printing techniques was investigated. Using the design rules outlined in Demir & Previtali [5] two stent designs were developed and the impact of etching as a post-processing technique on the stent performance was investigated. This was done using a combination of experimental testing combined with FE modelling. Further to this, using the CT reconstructed patient geometry, a FE model was developed to simulate the deployment of the stent into the patient’s anatomy and the performance of the stents pre- and post-etching were compared. To the author’s knowledge, this is the first time these methods have been applied to investigate 3D printed metallic stent design and performance.

## 2. Materials and Methods

### 2.1. Stent Design

Two stents were designed in Solidworks (Dassault Systemes Solidworks Corp, MA, USA) following rules of AM of stents based on Demir & Previtali [5]. The first stent design was a Simple Honeycomb (SH) stent design with a 10 mm diameter and 0.5 mm strut width and thickness shown in Fig. 2(A). The second stent design was a Novel Design (ND) using a honeycomb stent structure with a 10 mm diameter and 0.5 mm strut thickness, however, the width of the struts was varied around the circumference of the stent with all but two struts at each level of the stent having a width of 0.2 mm and two struts having a width of 0.5 mm as shown in Fig. 2(B).

**Figure 2.**
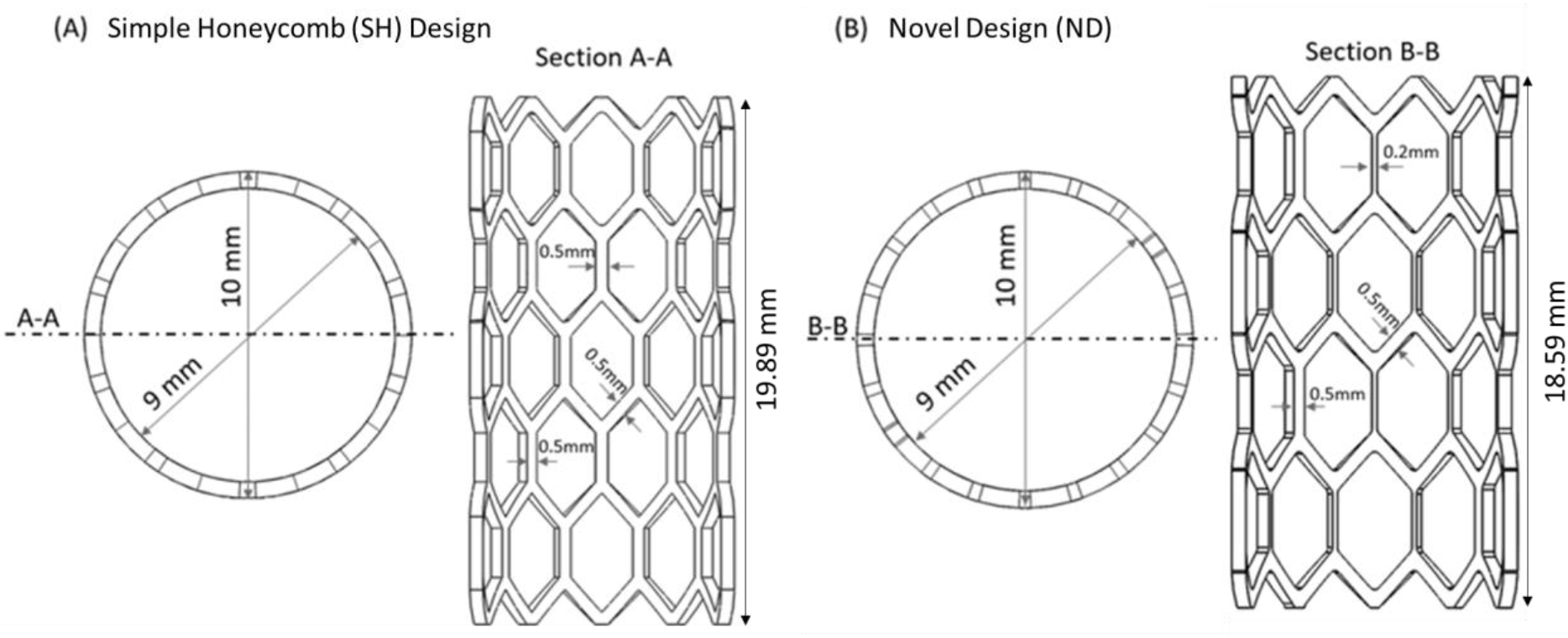
Schematic of (a) Simple Honeycomb (SH) and Novel Design (ND) stent designs.

### 2.2. Titanium Powder

The powder used was Ti-6Al-4V ELI (Grade 23) (Carpenter Additive, Cheshire, UK). The powder was spherical and produced by gas atomisation. Its size distribution was 22.5 - 47.2 μm with an average size of 32.7 μm. The powder’s chemical composition is given in Table 1:

**Table 1.**
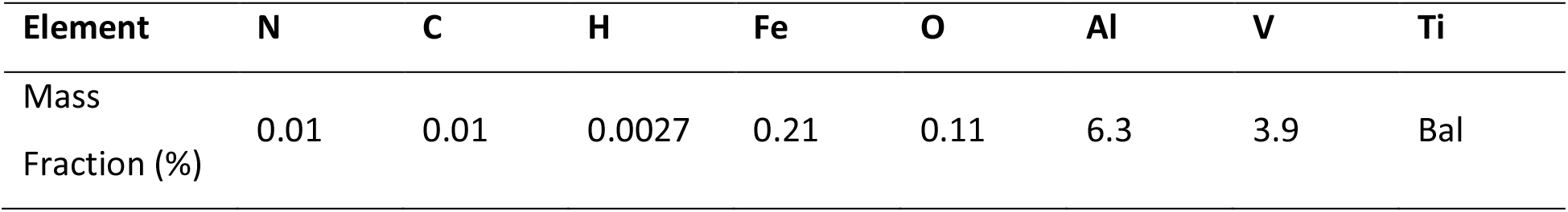
Chemical Composition of Titanium Powder

### 2.3. Selective Laser Melting

A ReaLizer SLM 50 with a 100 W fibre laser was used to print the stent geometries in Titanium. The estimated beam diameter was in the range of 15-20 μm. The processing chamber was filled with argon gas reaching an overpressure of 15 mbar. A circulation pump was used to maintain parallel flow to the powder bed and oxygen was maintained below 12%. A modulated continuous wave laser emission was used with an exposure time of 40 μs and a point distance of 10 μm. The laser current was maintained at 1200 mA for the outer boundary and 2800 mA for the inner hatching. A boundary-scan was executed first with parallel hatching scan lines with a hatch distance of 0.08 mm used internally. A layer thickness of 25 μm was kept constant throughout the build. Magics 23.02 (Materialise, Leuven, Belgium) was used to prepare the model. RDesigner (DMG MORI, Bielefeld, Germany) was used for setting laser trajectories and ROperator (DMG MORI, Bielefeld, Germany) was used for the machine preparation. Six of the SH stents and three of the ND stents designs were printed.

### 2.4. Post Processing (Etching)

Chemical etching efficiently reduces surface defects such as un-melted powder particles stuck to the built part and irregularities in layer stacking that manifests as protrusions perpendicular to the build direction [24]. The elimination of these defects leads to an increase in fatigue life when compared to as-built AM struts [25]. In this study, etching was used to reduce the stent strut thickness and improve the surface finish. The samples were immersed into an aqueous etching solution containing hydrofluoric and nitric acid ((HF: HNO3: H2O = 1:2:3) for 12 minutes and then transferred into distilled water for 5 minutes and rinsed with ethanol to complete the cleaning stage. Three of the SH design were etched, and all three of the ND designs were etched. The finished stent designs can be seen in Figure 3.

**Figure 3.**
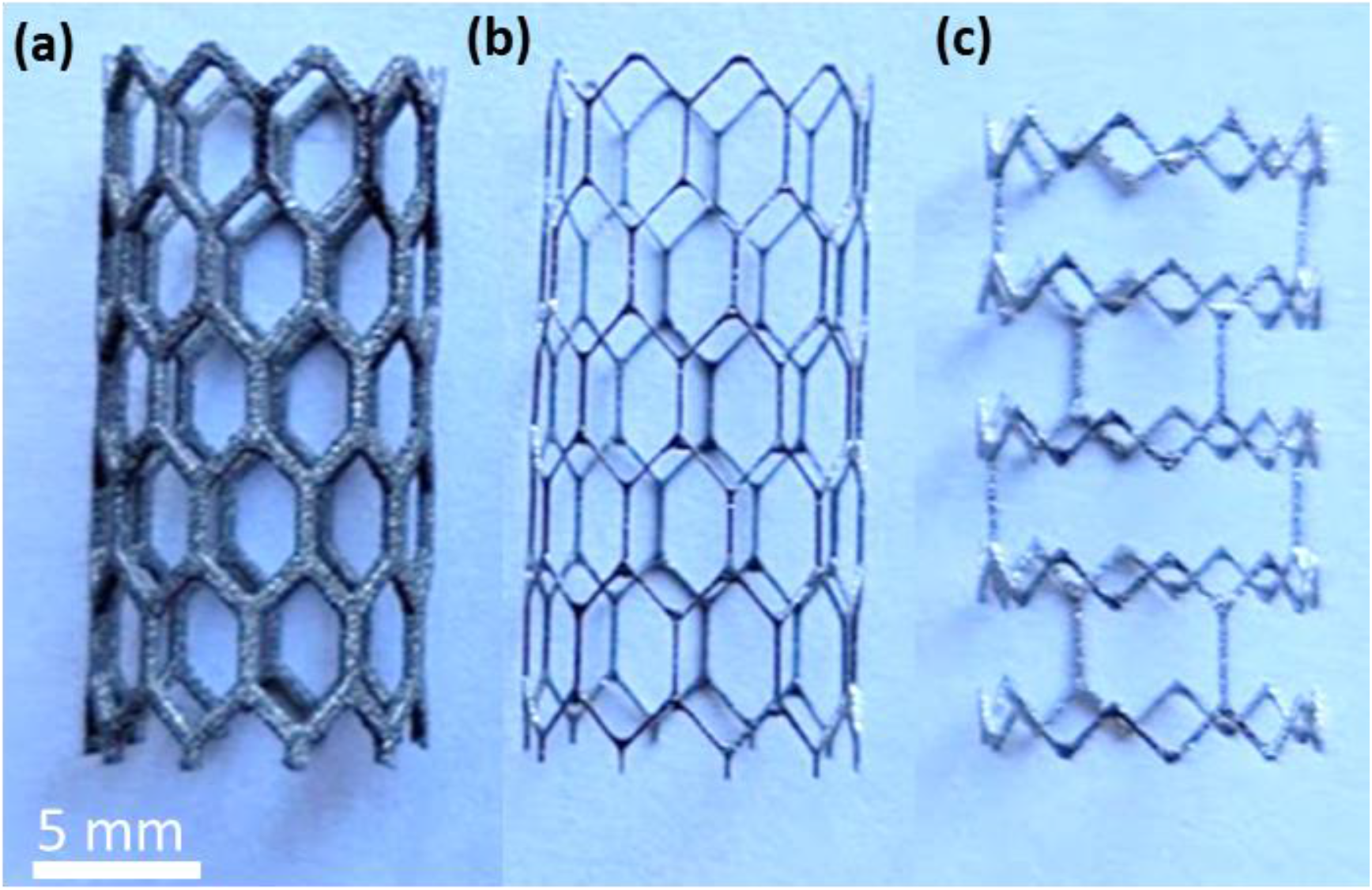
SLM Printed stents; (a) Simple Honeycomb (SH) (b) Simple Honeycomb Etched (SHE) and (c) the Novel Design Etched (NDE).

### 2.5. Printed Stent Geometric Dimensions

A stereoscopic microscope (Leica Microsystems, Wetzlar, Germany) was used to measure the width of the stents’ struts (Figure 4). Separate measurement of the vertical struts and 45° angled struts were taken. Struts thickness measurements were also taken. These measurements (Table 2) were then used to create representative geometries of each individual ‘as printed’ and ‘as etched’ stents using Solidworks (Dassault Systemes Solidworks Corp, MA, USA). The stents were then meshed using ANSA BETA (BETA CAE Systems, Luzern, Switzerland) and ABAQUS (SIMULIA, Providence, RI).

**Figure 4.**
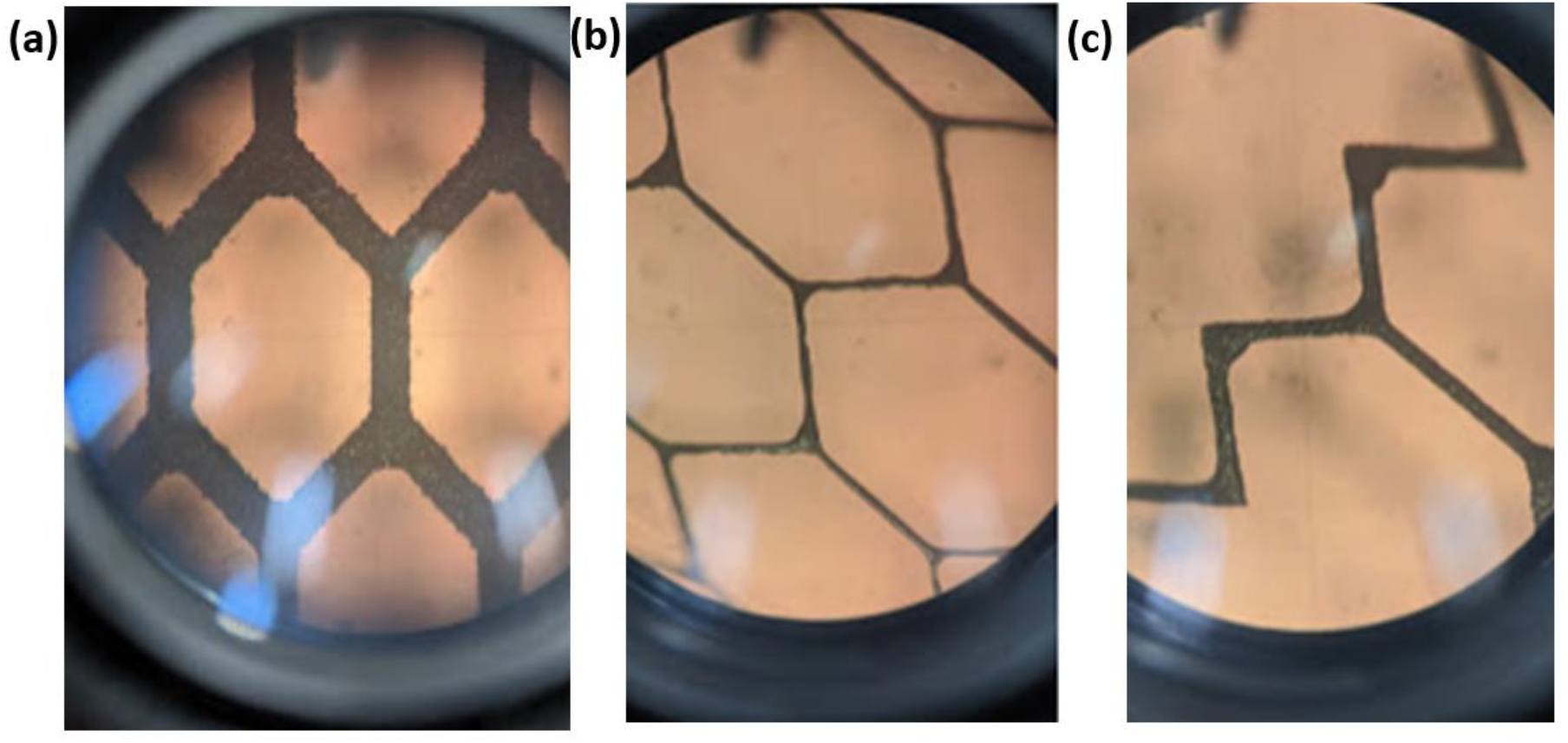
Stereoscopic images of stents’ struts of (a) Simple Honeycomb (SH) (b) Simple Honeycomb Etched (SHE) and (c) the Novel Design Etched (NDE) stent designs.

**Table 2.**
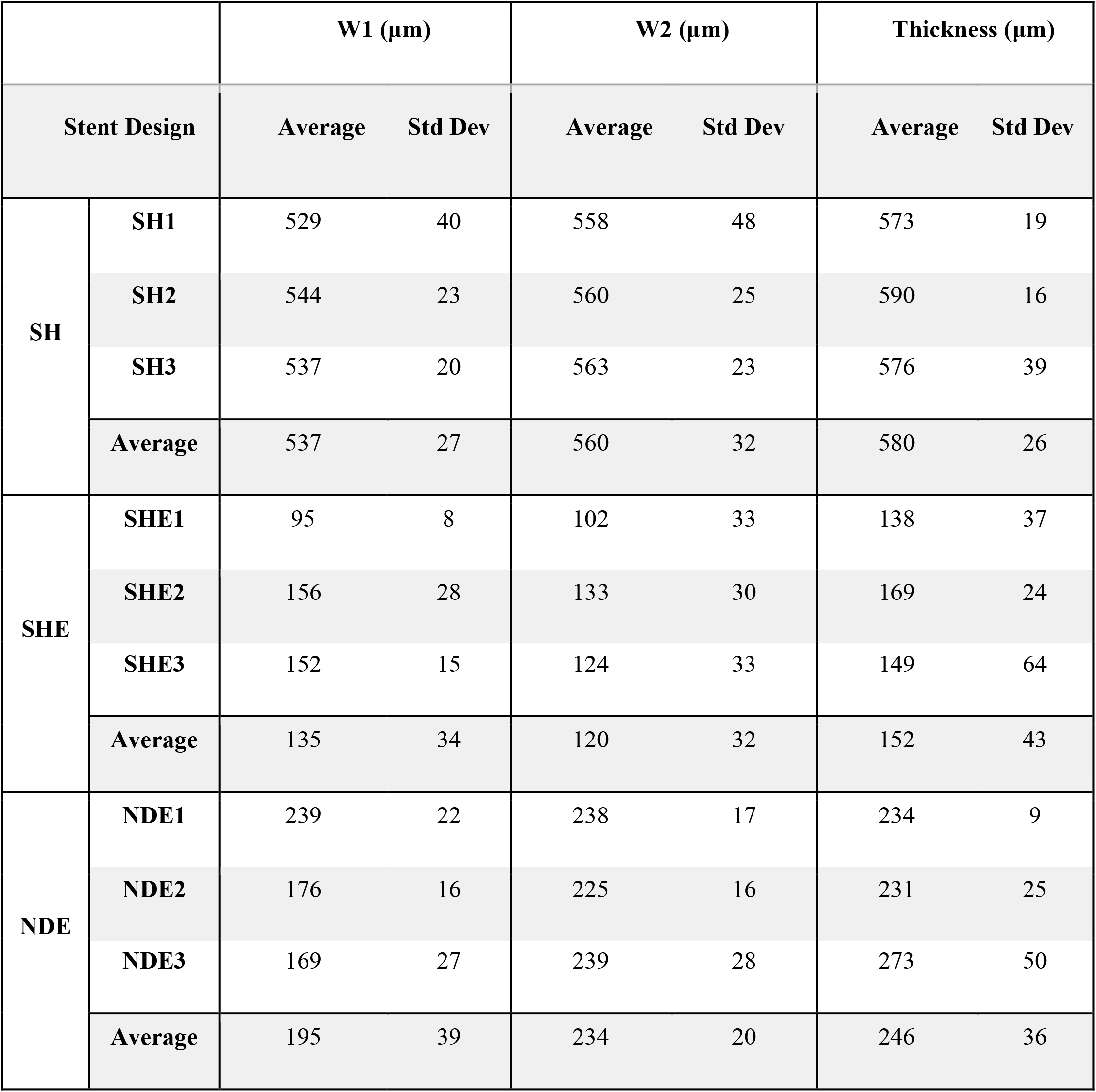
Stereoscopic measurements of stent vertical width (W1), width at the 45° (W2) and thickness.

|  |  | W1 (µm) |  | W2 (µm) |  | Thickness (µm) |  |
| --- | --- | --- | --- | --- | --- | --- | --- |
| Stent Design |  | Average | Std Dev | Average | Std Dev | Average | Std Dev |
| SH | SH1 | 529 | 40 | 558 | 48 | 573 | 19 |
|  | SH2 | 544 | 23 | 560 | 25 | 590 | 16 |
|  | SH3 | 537 | 20 | 563 | 23 | 576 | 39 |
|  | Average | 537 | 27 | 560 | 32 | 580 | 26 |
| SHE | SHE1 | 95 | 8 | 102 | 33 | 138 | 37 |
|  | SHE2 | 156 | 28 | 133 | 30 | 169 | 24 |
|  | SHE3 | 152 | 15 | 124 | 33 | 149 | 64 |
|  | Average | 135 | 34 | 120 | 32 | 152 | 43 |
| NDE | NDE1 | 239 | 22 | 238 | 17 | 234 | 9 |
|  | NDE2 | 176 | 16 | 225 | 16 | 231 | 25 |
|  | NDE3 | 169 | 27 | 239 | 28 | 273 | 50 |
|  | Average | 195 | 39 | 234 | 20 | 246 | 36 |

### 2.6. Experimental Crush Test

Using an Instron 3366 (Instron, MA, USA) a crush resistance test using parallel plates (Fig. 5) was performed on the three groups of stents; SH, SH-etched and the ND stent designs. The stent diameter was reduced by 3 mm and the reaction force was measured to quantify the stiffness of the different stents. A preload of 1.06 N was applied to each stent (0.56 N load as a result of the Perspex plate weighing 57.73g and 0.5 N preload applied by the machine).

**Figure 5.**
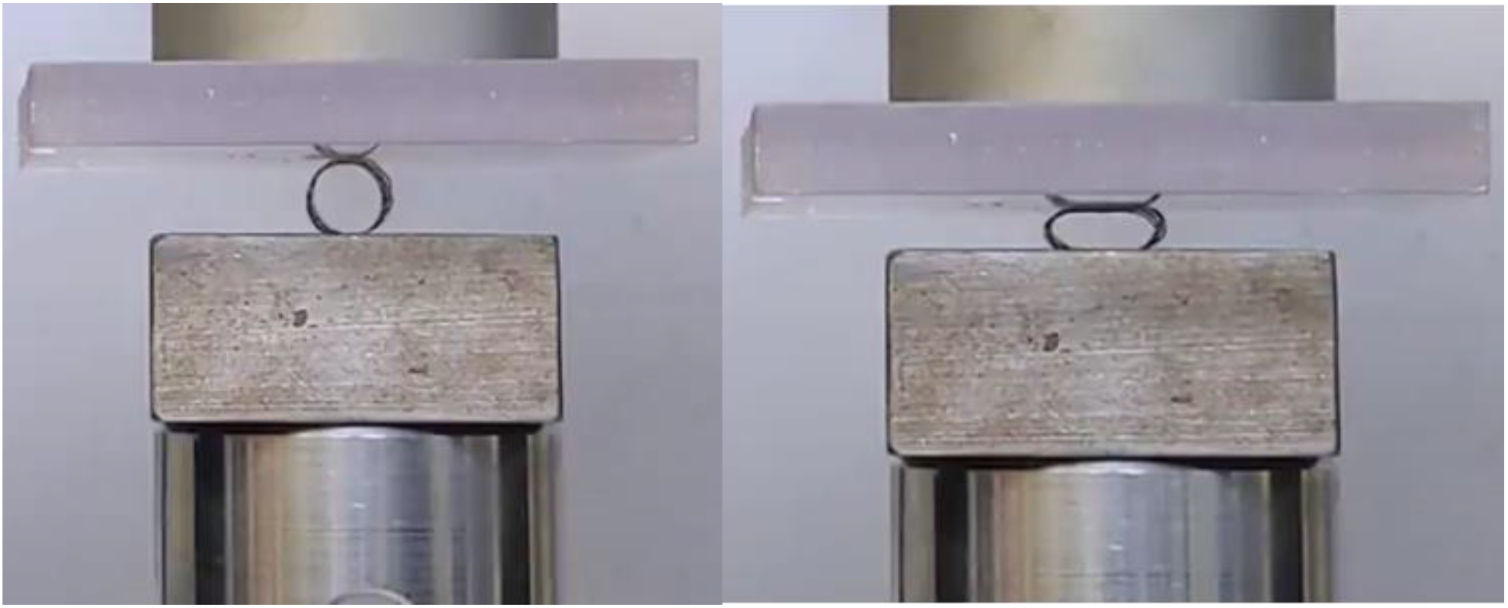
Experimental Parallel Plate Crush Test

### 2.7. Stent Surface Roughness

Scanning Electron Microscopy (SEM) images of the stents were taken after the crush test was performed to investigate the surface roughness of the stents. Additionally, White Light Interferometry (WLI) was used to assess the relative surface roughness of each of the stents to investigate differences between printed and etched stents and the variation of roughness between the strut directions within the stent designs. A ProFilm 3D (Filmetrics, Inc., San Diego, CA.) optical profilometer was used and arithmetical average roughness for a given path, R_a_, was used to compare surface quality.

ISO 4288:1996 [26] recommends a roughness sampling length of 0.8 mm and a roughness evaluation length of 4 mm for R_a_ values greater than 0.1 μm and less than or equal to 2 μm, corresponding to the R_a_ range for the etched stents. Similarly, the ISO standard [27] specifies a roughness sampling length of 2.5 mm and roughness evaluation length of 12.5 mm for R_a_ values greater than 2 μm and less than or equal to 10 μm, corresponding to the R_a_ range for the unetched stents. Due to limitations related to the length of the diagonal struts, it was not possible to attain this recommended sampling length. To mitigate this a consistent sample length of 600 μm was used across all measured struts and in a similar fashion to [27] and the effect of the low-test length was neglected as the measurements were used comparatively.

Eight struts, four vertical struts and four diagonal struts from each sample were scanned at a length of 600 μm. These images were obtained at 50x magnification, giving a field of view of 0.4 x 0.34 mm. Two scans were taken with a back-scan length of 110 μm and then stitched together with the provided profilometer software to achieve the prescribed sample length. The stitched images provided a 3D map of the surface topography, this was then processed within the software package to remove outliers whereby pixels with a slope above 60° were removed and invalid pixels were filled in. The line roughness tool was then used to calculate a R_a_ value along the strut within the scan length. Additional microscopy images at 10x and 50x magnification were taken for further qualitative comparison of surface roughness and defects.

### 2.8. Finite Element Crush Test

A FE model was created to replicate the experimental crush test using ABAQUS Standard 2017 (SIMULIA, Providence, RI). Two parallel plates 45 x 45 mm were meshed using (SFM3D4) 4-noded quadrilateral surface elements. The stent was placed between the two plates as seen in Fig 10(a). Zero friction and hard normal contact were assigned between the stent and the plates. The bottom plate was fixed, and the top plate was displaced −3 mm in the y-direction to replicate the displacement in the crush test. The material properties used were linear elastic with a Young’s modulus that was determined using the inverse FE approach outlined in Section 2.9 and a Poisson’s ratio of 0.31[28,29]. This test was used to test the SH design, SH-etched design and the ND stents. A mesh dependency was performed on each variation of the stent design whereby a 100% increase in mesh density led to < 2% change in the final reaction force.

### 2.9. Inverse Finite Element Analysis

Calibration of the Young’s modulus for each of the stents was carried out using an optimisation loop in MATLAB (MathWorks, MA, USA). An initial estimate of the Young’s modulus for each stent was made based on the experimental crush tests. This value was then used in the crush test simulation and the force-displacement results were then compared to the experimental data in the linear elastic region by calculating the Root Mean Square Error (RMSE). This process was repeated until the solver reached the lower bound tolerance of 1e^-4^ for the RMSE value.

### 2.10. Patient-Specific Finite Element Model of Stent Deployment

Fully anonymised CT scan data of the aortic arch of a paediatric patient with an aortic coarctation, necessitating the placement of a stent, were obtained from our clinical collaborator. Image segmentation was applied to this CT data to extract a computer-aided design (CAD) representation of the patient geometry using 3D Slicer [30]. A skin surface of the vessel was then constructed with ANSYS SpaceClaim (2019 ANSYS. Inc) and finally, a volumetric hexahedral mesh representation of the geometry was generated with ANSA BETA Pre-Processor (BETA CAE Systems, Luzern, Switzerland) (Fig 6). The aortic arch was assigned a uniform thickness of 1 mm [31] and was meshed using (C3D8R) 8-node linear brick elements with reduced integration and hourglass control. A mesh dependency analysis was performed on the aortic arch and it was found there was a 1.9% change in the max von Mises stress and a 2.4% change in the average von Mises stress with a 59% increase in the mesh density.

**Figure 6.**
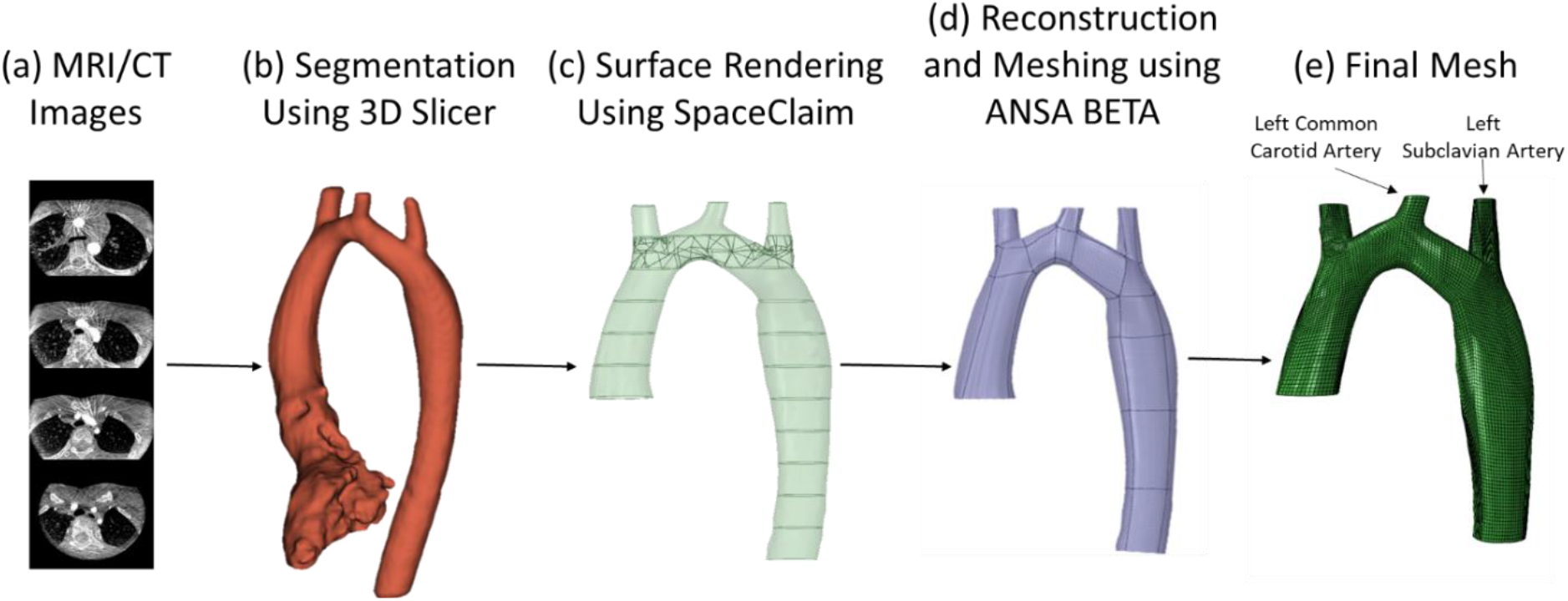
Steps used to generate a patient-specific finite element mesh for CT data of a patient. (a) CT/MRI images (b) segmentation using 3D Slicer (c) surface rendering using SpaceClaim (d) reconstruction and meshing using ANSA BETA (e) final meshed geometry.

Using ABAQUS Explicit a finite element model was used to simulate the deployment of the three stents into the patient’s anatomy. The stents were modelled as linear elastic titanium and meshed as in the FE crush test. The patient’s aortic arch was modelled as an incompressible hyperelastic isotropic Mooney–Rivlin model. The model was defined using the following strain energy density function, U:

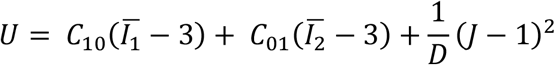

where *C*_10_ and *C*_01_ are material parameters, D is the incompressibility parameter and *I*_1_ and *I*_2_ are the first and second invariant of the right Cauchy–Green deformation tensor, ***C***:

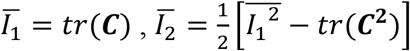

D was assigned a value of 1e^-6^ assuming near incompressibility [32,33]. The material constants *C*_10_= 0.5516 MPa and *C*_01_ = 0.1379 MPa were adopted from [33–35].

Using a cylindrical part (crimper) meshed using (SFM3D4R) 4-noded quadrilateral surface elements with reduced integration and a local cylindrical coordinate system was created to displace the crimper radially inward. Hard normal contact with a coefficient of friction of 0.8 was applied between the stent and the crimper [36]. The crimper was assigned as the master surface, and the frame was assigned as the slave surface.

The top and bottom edges of the aortic arch were constrained using displacement boundary conditions in the longitudinal and circumferential directions. The stent was uncrimped and hard normal, frictionless contact was applied between the stent and the aorta [37]. The kinetic energy of the simulations was monitored to ensure that the ratio of kinetic energy to internal energy remained less than 5%. This criterion was proposed by [38] and ensures that the dynamic effects are negligible.

The patient-specific model was used to compare the performance of the all three stent designs and examine the material properties that were calibrated using the inverse FE approach.

## 3. Results

### 3.1. As Printed and As Etched Stent Geometries

#### 3.1.1 Geometric Differences

Stereoscopic measurements of the as printed SH stent geometries found that stents had increased width and thickness compared to the STL input. It was found that the thickness of the stents showed the greatest increase, averaging 80 ± 26 μm. In struts printed at 45°, an average increase of 60 ± 32 μm in width was observed compared to an average increase of 37 ± 27 μm in width in struts printed vertically.

Etching was found to successfully remove material from the width and thickness of the stents leading to an average thickness of 152 ± 43 μm for the SHE and 246 ± 36 μm for the NDE stents. The average width of struts at 45° was found to be 120 ± 32 μm for the SHE and 234 ± 20 μm for the NDE while the vertical strut widths were found to be 135 ± 34 μm and 195 ± 39 μm for the SHE and NDE respectively. Furthermore, the 200 μm struts present in the NDE stent prior to etching were completely removed as a result of etching. This led to an open-cell stent design that to the author’s knowledge has not previously been achieved through 3D printing.

#### 3.1.2 Surface Roughness

In addition to the removal of material from the thickness and width of the stent, etching led to an improvement in surface roughness. Figure 7 demonstrates an SEM image of a stent pre- and post-etching where the improvement in the surface finish after etching is evident. Un-melted powder particles and irregularities in layer stacking were successfully removed using chemical etching upon visual inspection.

**Figure 7.**
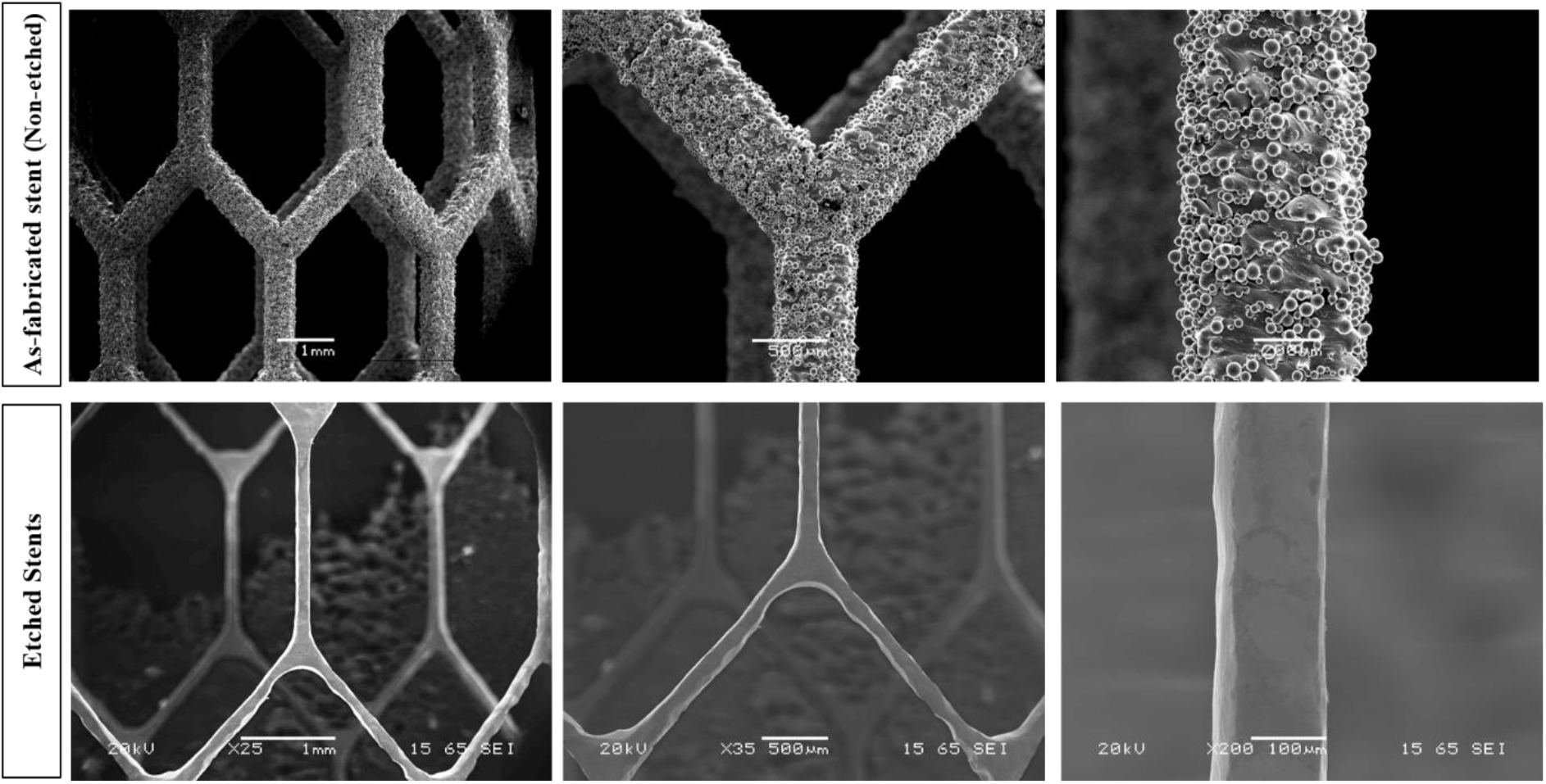
SEM micrographs of Honeycomb stents, showing the effects of chemical etching on surface cleaning and reduction of strut sizes. In as-fabricated status (top row) and after etching (bottom row).

WLI was performed to quantify the improvement in surface roughness evident from SEM images and investigate the relationship between print angle and surface roughness. A box and whisker plot of the arithmetical average roughness, R_a_, for each stent group is presented in Figure 8(a). In terms of mean and standard deviation the R_a_ values for the SH, NDE and SHE were measured as 3.76 ± 0.27 μm, 1.13 ± 0.14 μm and 0.39 ± 0.06 μm respectively. To investigate statistically significant differences between the means of each group a Welch’s t-test with a 95% confidence interval was applied. This test method assumes the means of the compared populations are normally distributed and is insensitive to equality of variance. The t-tests estimate with 95% confidence that there is a statistically significant difference between each of the stent groups.

**Figure 8.**
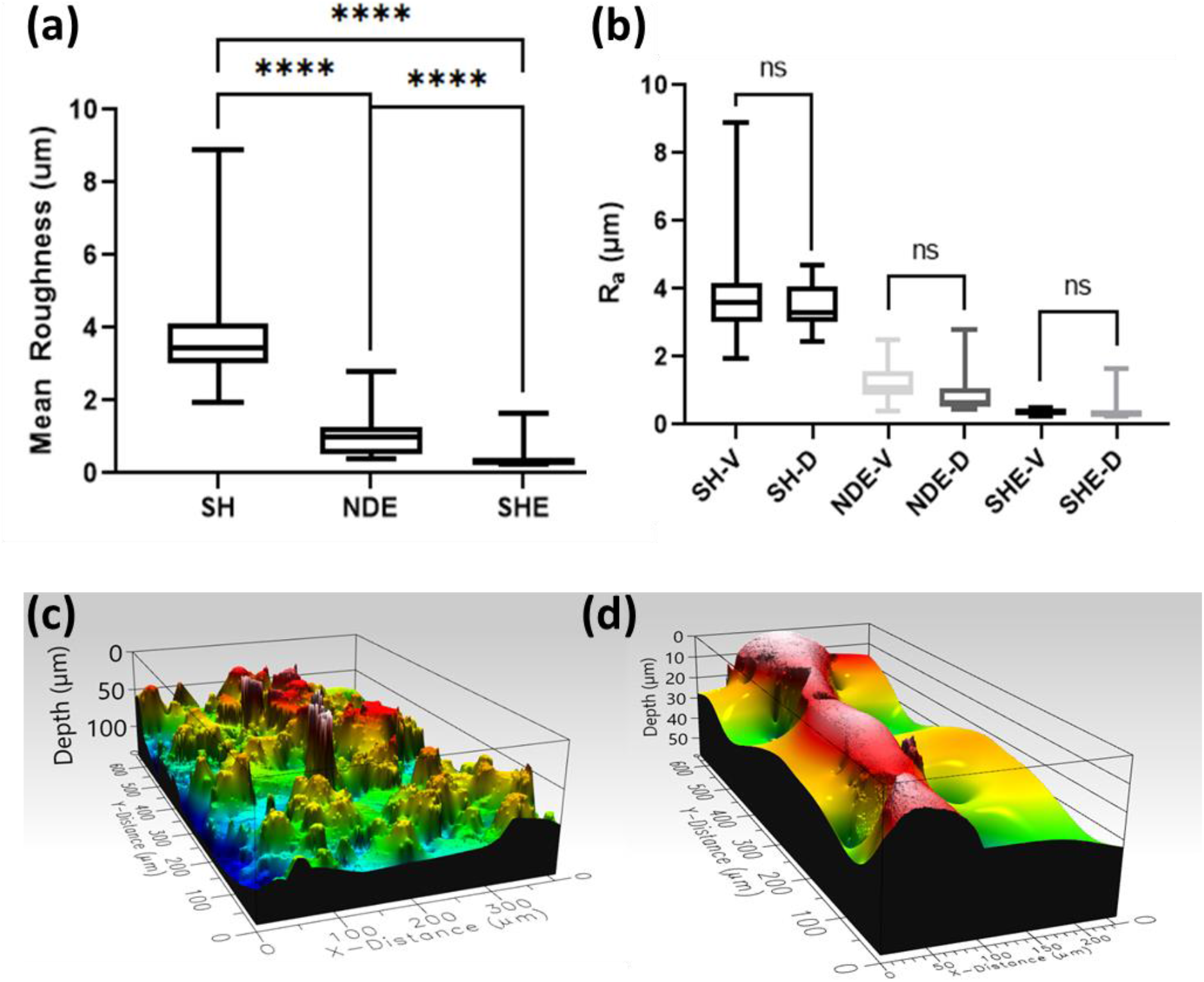
(a) Box and whisker plot of arithmetical average surface roughness, R_a_ (μm) and results of Welch’s t-test for Simple Honeycomb (SH), Simple Honeycomb Etched (SHE) and the Novel Design Etched (NDE). **** indicates a p-value less than or equal to 0.0001. (b) Box and whisker plot of arithmetical average surface roughness R_a_ (μm) and results of Welch’s t-test for vertical (V) and diagonal (D) strut directions for each stent group. ns indicates no statistically significant difference between means. (c) WLI 3D surface map of an SH stent strut and (d) SHE stent strut.

Figure 8(b) presents a box and whisker plot of the R_a_ value of each stent group separated by the two strut directions, namely the vertical (V) and diagonal (D) directions. A Welch’s t-test was again applied to investigate statistically significant differences between horizontal and diagonal struts within the stent groups. The analysis estimates with 95% confidence that there is no statistical difference between the surface roughness for different strut directions within stent groups.

### 3.2. Experimental Crush Test

The results of the experimental crush test are shown in Figure 9. The results demonstrate the influence that etching has on stent performance. It must be noted that there was a broken strut observed in SHE-1 prior to the experimental crush test. It must also be noted that due to a lack of symmetry in the NDE stents greater variation is seen in the crush test results when compared to the SH and SHE stent designs.

**Figure 9.**
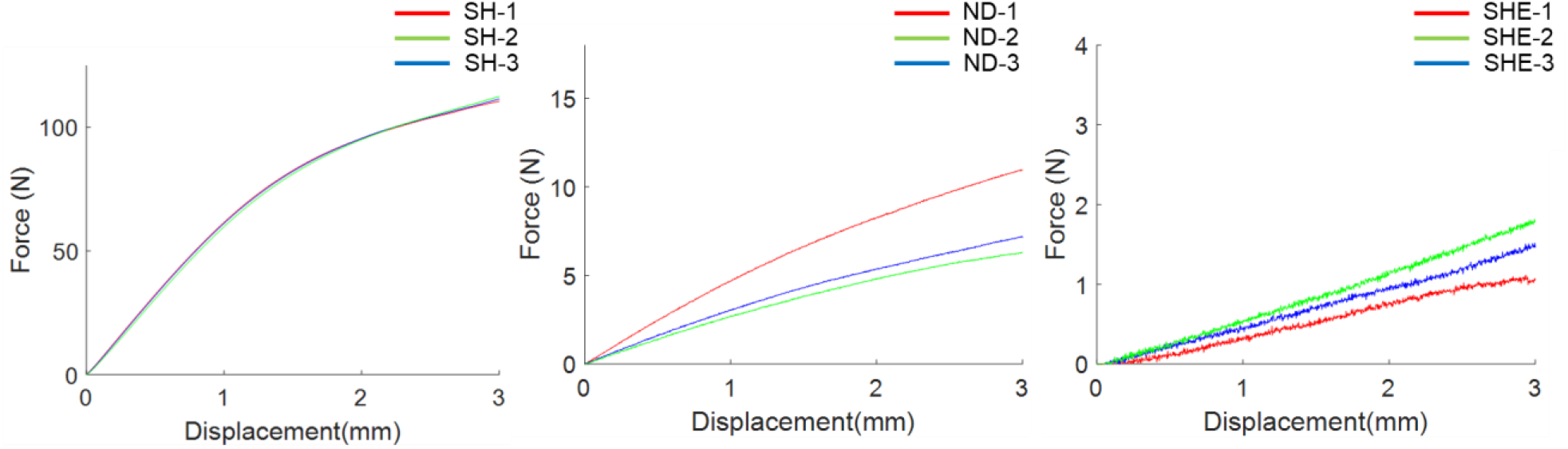
Force-displacement curve from the experimental crush test of the printed stents.

### 3.3. Finite Element Crush Test and Material Calibration

Figure 10 (b), (c) and (d) shows the force displacement curves of the experimental crush test to the corresponding FE prediction using the calibrated Young’s modulus for the different stent designs. Table 3 demonstrates the Young’s modulus and corresponding RMSE values calculated using the inverse FE approach outline in Section 2.9. The SH designs show a decreased stiffness compared to the SHE and NDE stents. The average Young’s modulus (E) for the SH stent was found to be 33.89 GPa when compared to an average of 76.84 GPa for the SHE stent designs. The SHE-1 design was not included due to the fact that the broken strut could not be accurately represented in an FE model and would have inaccurately influenced the material characterisation.

**Figure 10.**
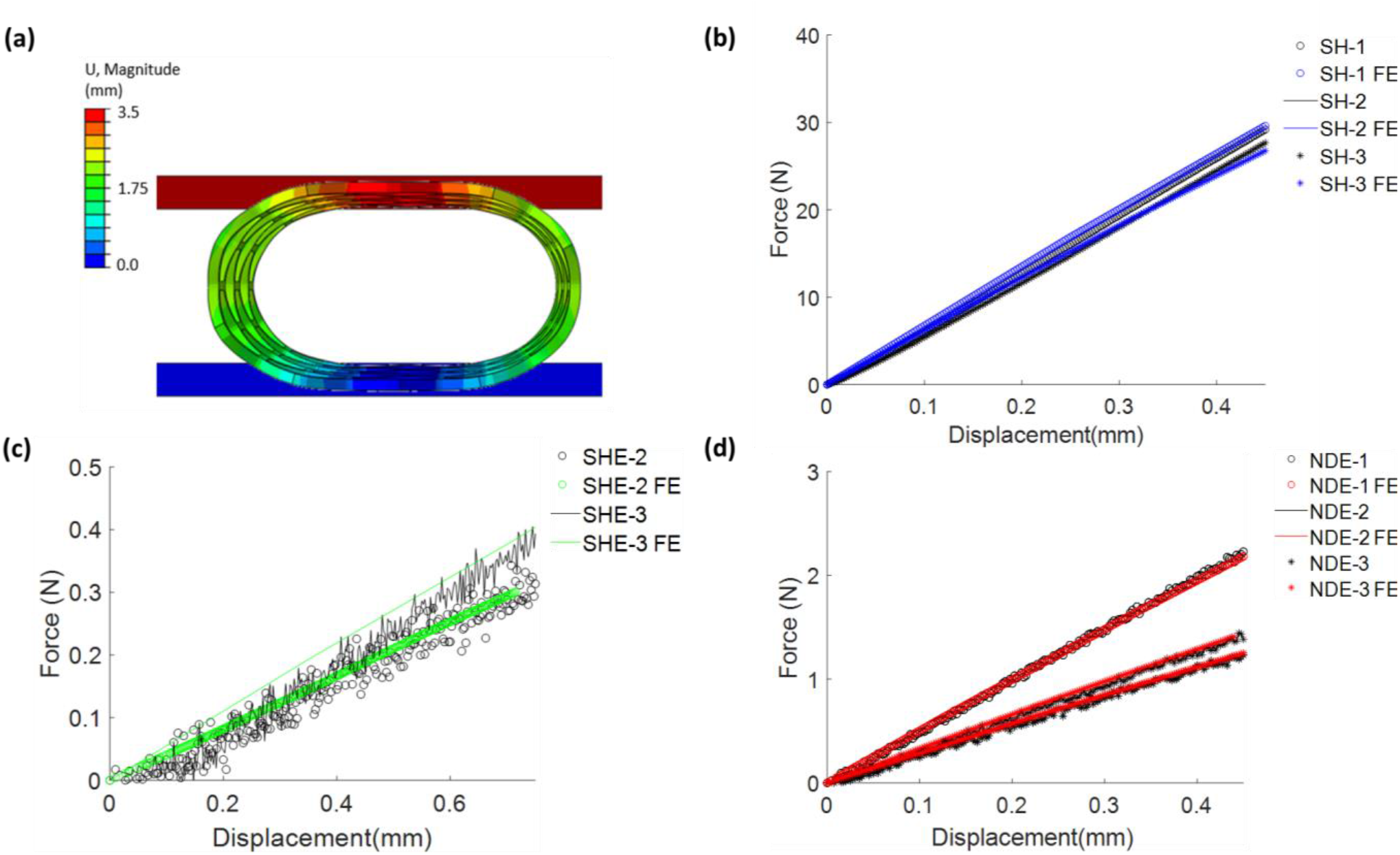
(a) Contour plot of displacement for the finite element model of the crush test of SH stent. Force-displacement curve from the experimental crush test results and finite element (FE) predictions for (b) SH (c) SHE and (d) NDE stent designs.

**Table 3.** Calibrated Young’s Modulus and corresponding RMSE value for each stent design.

| **Stent Design** | **Youngs Modulus (GPa)** | **RMSE** |
| --- | --- | --- |
| SH1 | 35.794 | 0.447 |
| SH2 | 33.809 | 0.576 |
| SH3 | 31.059 | 0.703 |
| SHE2 | 55.018 | 0.024 |
| SHE3 | 98.655 | 0.284 |
| NDE1 | 110.480 | 0.042 |
| NDE2 | 114.042 | 0.020 |
| NDE3 | 63.957 | 0.023 |

### 3.4. Patient-Specific Finite Element Model of Stent Deployment

The patient-specific model was used to compare the performance of the three stent designs and examine the material properties that were calibrated using the inverse FE approach. Six designs were investigated and the average Youngs moduli from the SHE (76.84 GPa) and SH (33.89 GPa) stent designs were examine using the model. The models investigated the SH-as printed (E=33.89 GPa), SH-as printed (E=76.84 GPa), SH-STL (E=33.89 GPa), SH-STL (E=76.84 GPa), SHE (E=76.84 GPa) and NDE (E=76.84 GPa) stent designs. Stent performance was investigated in terms of stress in the aorta and lumen gain. Figure 11 displays contour plots of von Mises stress in the aortic arch for each different stent design. It can be seen that the NDE and SHE stents results in lower stress distributions in the aortic arch than the SH stent designs.

**Figure 11.**
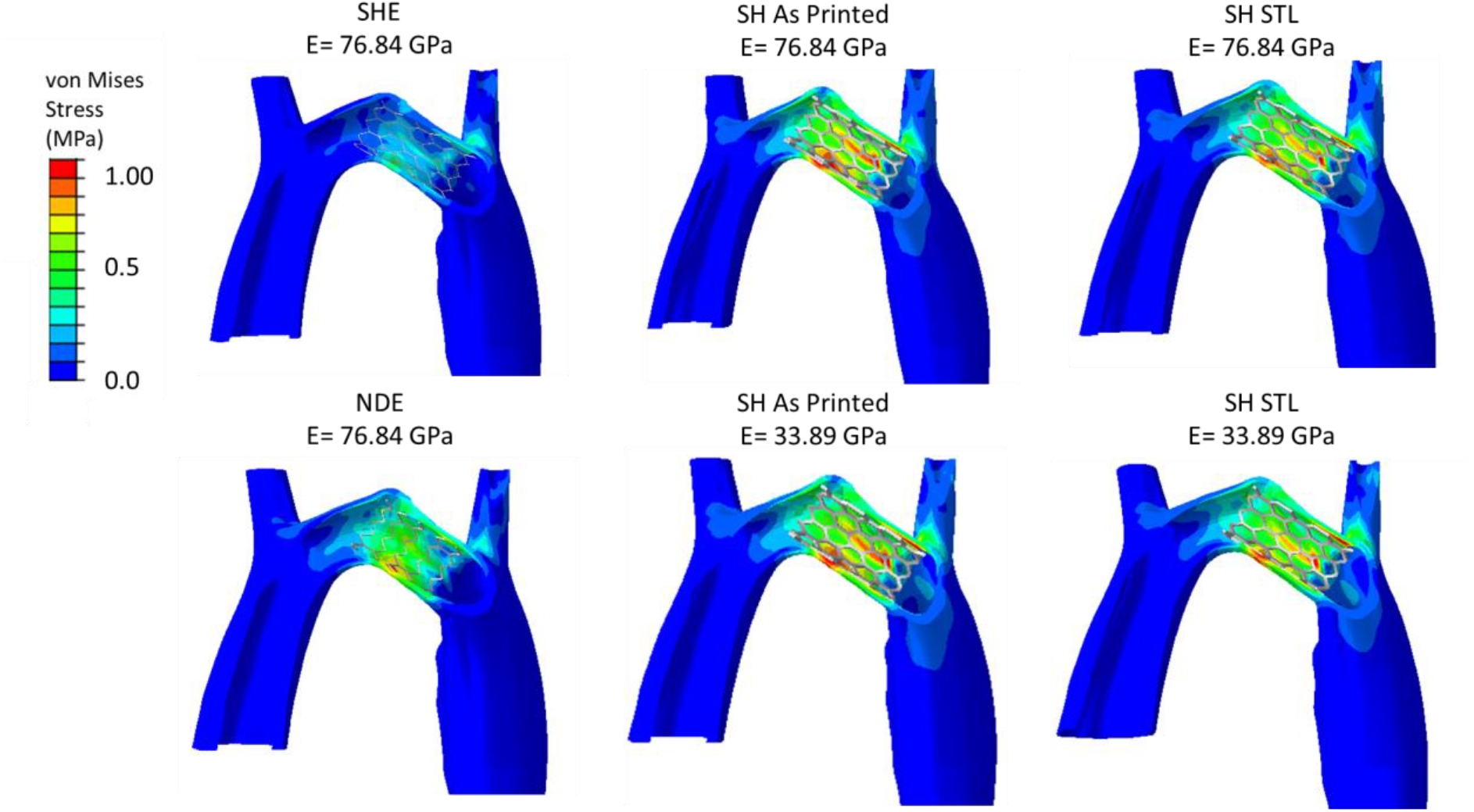
Contour plots of von Mises stress in the aorta post-deployment of different stent designs.

The section of the aortic arch between the left subclavian artery and the left common carotid artery was examined to investigate lumen gain and maximum von Mises stress (Table 4). It was found that while the SHE design reduced stress it also led to a lower increase in lumen area. Further to this, it was found SH ‘as print’ design led to the greatest increase in maximum von Mises stress and the greatest increase in lumen area.

**Table 4.**
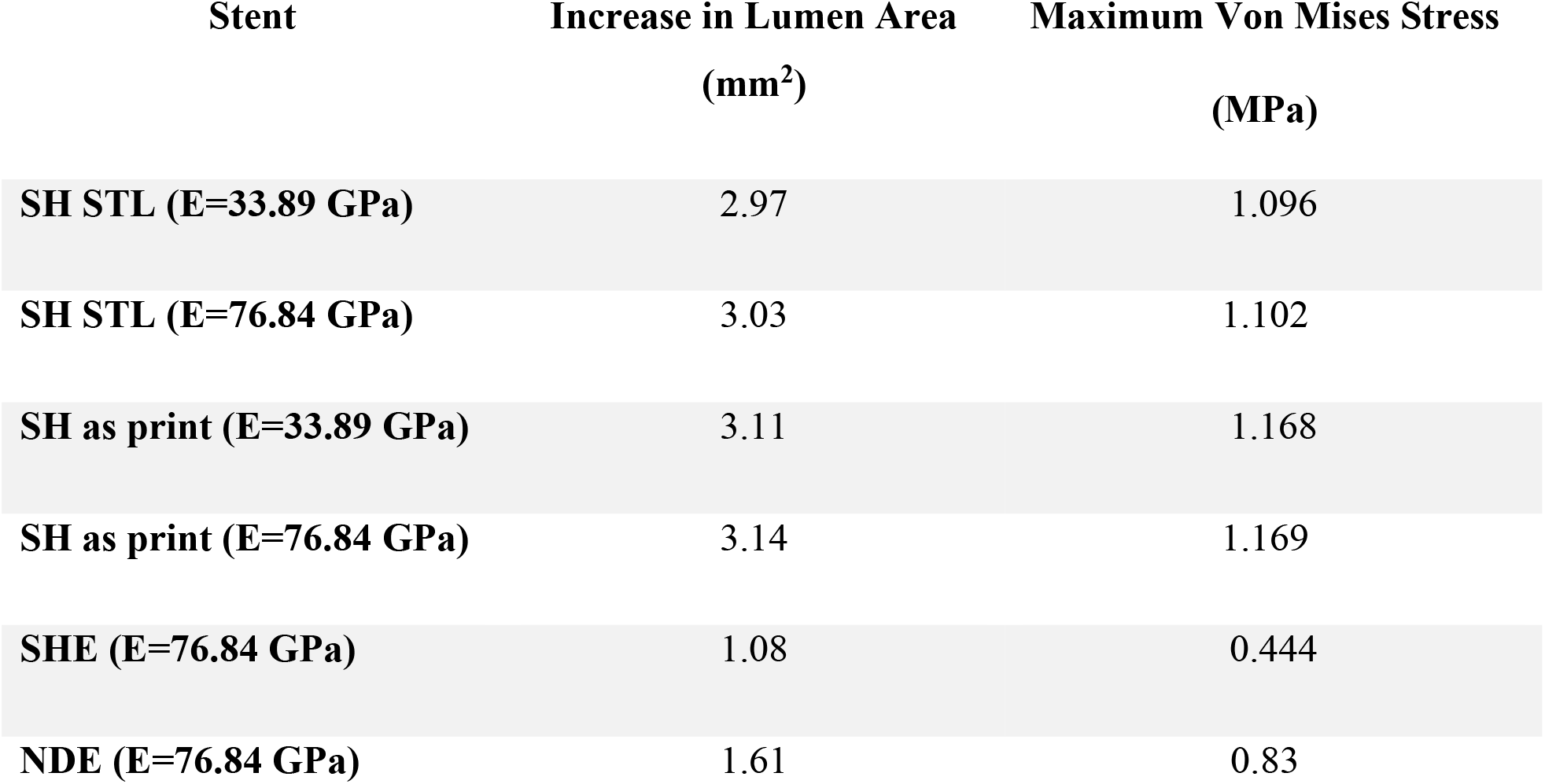
Maximum von Mises stress and Increase in Lumen Area for the different stent designs and stiffnesses.

| **Stent** | **Increase in Lumen Area**<br>**(mm<sup>2</sup>)** | **Maximum Von Mises Stress**<br>**(MPa)** |
| --- | --- | --- |
| **SH STL (E=33.89 GPa)** | 2.97 | 1.096 |
| **SH STL (E=76.84 GPa)** | 3.03 | 1.102 |
| **SH as print (E=33.89 GPa)** | 3.11 | 1.168 |
| **SH as print (E=76.84 GPa)** | 3.14 | 1.169 |
| **SHE (E=76.84 GPa)** | 1.08 | 0.444 |
| **NDE (E=76.84 GPa)** | 1.61 | 0.83 |

Finally, a percentage volume plot demonstrating the percentage volume of tissue at different bands of von Mises stress was used to examine the stress distribution in the tissue between the left subclavian artery and the left common carotid artery (Figure 12). It was found the SHE and NDE designs led to a considerable reduction in the stresses observed in the tissue.

**Figure 12.**
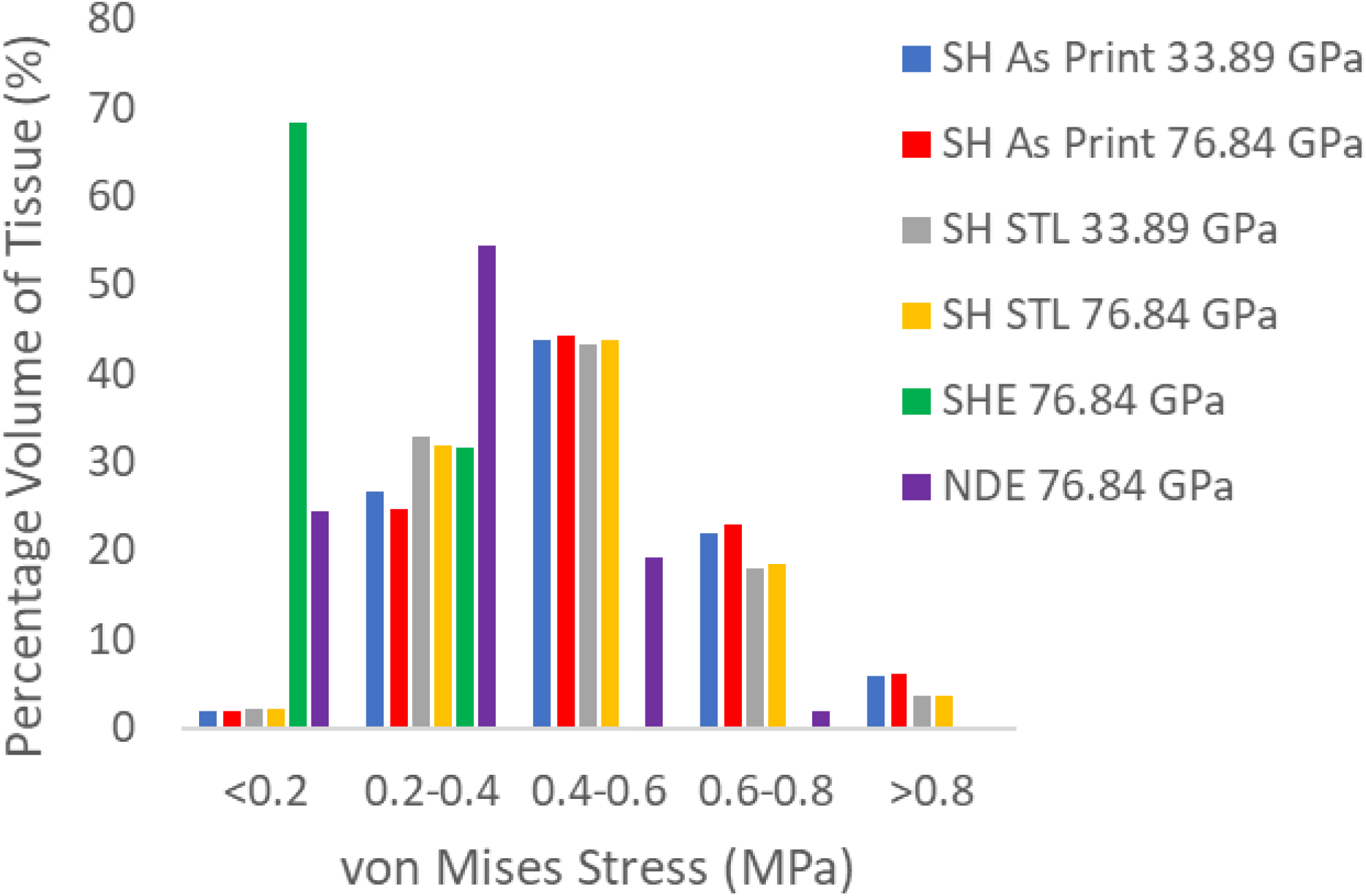
Percentage of volume graph for the different stent designs and stiffnesses showing the stress distribution between the left subclavian artery and the left common carotid artery.

## 4. Discussion

In this study, we have investigated the use of SLM printing to design patient-specific stents. Using FE methods, the performance of these stents was assessed alongside experimental bench tests. FE methods were further used to compare the performance of the stents post-deployment in the patient’s anatomy. Although there is still much to be investigated for the full potential of SLM printed stents to be realised, this study demonstrates the suitability of the methods outlined for use in the design of SLM printed stents for patient-specific purposes, in particular, in relation to paediatric patients. The results of this study demonstrate the ability of post-processing techniques such as etching to overcome design limitations that are inherent to the SLM technique and to better control the stent designs available through AM.

Several limitations and assumptions have been made throughout this study, firstly the properties for titanium used in this model are linear elastic material properties however, the stents would likely undergo plastic deformation during stent deployment. It must be noted that the material calibration was used to highlight how overestimation of stent dimensions due to melted powder particles and irregularities in layer stacking can lead to different outcomes and this was adequately achieved despite the use of a linear elastic approach. Similarly, the aortic arch was modelled as a hyperelastic isotropic material although it is known that arterial tissue exhibits anisotropic behaviour [39]. However, the constitutive laws used were fitted to the circumferential direction, the stiffest directions of the aortic root, and it is known that the aortic root responds to stent deployment predominantly in the circumferential direction. Therefore, the use of an isotropic model was considered to be appropriate for correctly capturing the stress in this model [33,40,41]. Moreover, this model was only used for comparison purposes of the stent designs. It must also be noted the stents modelled were taken as approximate measurements of the thickness and width of the stents and the variations in measurement observed throughout the stent were not modelled. However, this approximation was deemed valid as several measurements were taken from different struts throughout each stent geometry and microscopic measurements were deemed preferable to CT due to concerns over beam hardening and blooming effects when taking CT of titanium [42]. Finally, it must be noted that the Ti-64 was used predominantly for ‘proof-of-concept’ purposes and future work will include the investigation of other metal powders, such as Nitinol, to optimize cardiovascular stent designs.

One of the main findings of this study was a novel method to overcome some of the constraints in SLM printing where stents need to be self-supporting. Using thinner struts that can be etched away can generate a different design and create new capabilities for 3D printed stents, even allowing the creation of open-cell designs. Open-cell designs can be advantageous in reducing obstruction to branching arteries and can better conform to tortuous arteries as they have superior flexibility to close cell designs [43]. It must be noted that although only a crush test was carried out to assess stent performance in this study, future work will include a more exhaustive analysis of stent performance and include 3-point bend testing to more quantitively assess the stent bending stiffness.

Further to this, using etching, thinner strut widths and thicknesses can be achieved which can lead to a reduction in stiffness of the overall stent performance, as shown in the experimental crush tests. The strut thickness that can be achieved with SLM printing is dependent on several limiting factors including laser spot size and resolution [5,13], and using etching can further reduce the thickness of struts to levels that cannot be successfully achieved during printing.

Furthermore, etching was found to significantly improve stent surface roughness which can improve fatigue life [25]. A reduction in surface roughness of 3.37 μm was achieved between the SH and the SHE stents, where SHE stents had a surface roughness of 0.39 ± 0.06 μm. This is a distinct improvement on previously published R_a_ values of 2.2 μm measured post electropolishing in a recent study [44]. Surface roughness values for SHE and NDE stent designs were also statistically significantly different from one another. It must be noted that the SHE stents and NDE stents were etched using the same procedure. However, the two stent designs were etched on different days using different batches of etchant. This raises questions regarding the repeatability of the current etching procedure and future work will include a more robust etching study examining the repeatability and reproducibility of the etching protocol. Furthermore, etching led to variation in the width and thickness throughout individual struts. Future work will also include examining methods to improve the consistency of material removal throughout a stent.

The experimental crush test demonstrates a clear reduction in the overall radial stiffness of the stent post-etching. However, the material calibration of the reconstructed ‘as printed’ and ‘as etched’ stent designs found the material in the SH (non-etched) stents to be more compliant. In fact, both the SHE and SH stents were printed at the same time under the same conditions and print parameters. The reason for this perceived increase in compliance can be attributed to melted powder particles and irregularities in layer stacking contributing to increased stent dimensions but not contributing to the stent’s load-bearing ability and the overall stent performance. It must also be noted when examining the RMSE values for the SH stent, that the values achieved were higher than those observed for the SHE and NDE stents. This further supports the argument that the dimensions taken from these stents do not accurately represent the amount of load-bearing material. Such discrepancies in the amount of load-bearing material must be carefully considered when predicting SLM stent performance.

Finally, the stent performance was tested in a patient-specific FE model. This was used to assess the difference in the performance of the different stent designs and different material properties of the stents. It was found that the SH ‘as printed’ gave the greatest lumen area gain which can be attributed to the added thickness. However, it also led to the highest stresses in the aortic root. In contrast, the SHE stents design lead to the lowest gain in lumen area with the lowest stress experience in the aortic root. The advantage in the NDE design can be seen in the reduction of occlusion to the branching left common carotid artery in Figure 11. Future work will combine such FE models with optimization tools to develop an optimal stent design that can reduce occlusion to branches and maximise lumen gain while minimising stresses in the aortic arch.

## 5. Conclusion

SLM printing of stents is still an ongoing area of research with continued investigation needed to realise its full potential. This study demonstrates the potential for etching techniques to overcome some of the challenges faced when designing SLM printed stents. Further to this, FE modelling has great potential to both inform these designs and enable in silico testing of designs for specific patients. Future work aims to apply the framework developed in this paper to optimise patient-specific paediatric stent designs that can be printed using the SLM technique.

## 6. Acknowledgements

This study was funded National Children’s Research Centre, Crumlin (Grant No. G/19/3), an Irish Research Council Government of Ireland Postdoctoral Fellowship 2019 (GOIPD/2019/222) and from research supported in part by a research grant from Science Foundation Ireland (SFI) under the grant number 12/RC/2278_2. Prof. Moataz M. Attallah and Dr Parastoo Jamshidi acknowledge the support from Engineering and Physical Science Research Council (EPSRC) (Grant Number: EP/R001650/1; Title: Smart peripheral stents for the lower extremity–design, manufacturing and evaluation).

## Notes

### Competing Interest Statement

The authors have declared no competing interest.

